# A shared binding interface controls Vps13 organelle-specific targeting independently of its vacuolar protein sorting function

**DOI:** 10.1101/2025.08.12.669934

**Authors:** Kevin Ryan Jeffers, Samantha Katarzyna Dziurdzik, Michael Davey, Elizabeth Conibear

## Abstract

Yeast vacuolar protein sorting 13 (Vps13) is a bridge-like transporter that directs lipid flow between membranes at organelle contact sites. Vps13 targeting relies on organelle-specific adaptors containing proline-X-proline (PxP) motifs, which compete for binding to the Vps13 adaptor-binding (VAB) domain. Though a VAB-PxP interface has been identified for the mitochondrial adaptor Mcp1, whether other adaptors use identical binding mechanisms is unknown. Moreover, not every Vps13 function is connected to a known PxP adaptor, suggesting other adaptors may exist.

Here, we validate the significance of the shared VAB-PxP interface by showing that mutations within this region inhibit both adaptor binding and Vps13 membrane targeting in vivo. Using predictive modeling, we demonstrate that while adaptors share a common Vps13-binding interface, slight differences between these interfaces may contribute to preferential binding and adaptor competition. Notably, we find that the VPS pathway functions independently of the PxP motif binding site, and extensive scanning mutagenesis did not reveal a dedicated VPS-specific adaptor interface. Our results indicate the VPS pathway likely employs a non-PxP adaptor mechanism, yet the structural integrity of the VAB domain remains essential for proper pathway function.

**Significance statement:**

- Adaptor-mediated recruitment of yeast Vps13 to organelles requires a consensus PxP motif. It is unclear if all adaptors bind Vps13 in a similar manner and if all Vps13 functions are dependent on adaptors that contain this motif.
- We show that Vps13 binds the core PxP motifs from known adaptors through a shared interface, yet this interface is dispensable for Vps13’s role in the vacuole protein sorting pathway.
- Our work suggests that Vps13 engages in a wider range of targeting mechanisms than was previously recognized, and that the Vps13 adaptor-binding domain must have a function beyond targeting this consensus motif.

## INTRODUCTION

Interorganellar lipid trafficking is essential for cellular growth and function (Jackson *et al*., 2016; Wong *et al*., 2019). At membrane contact sites, specialized transporters orchestrate the movement of lipids between organelles. Vacuolar protein sorting 13 (Vps13) belongs to a family of large bridge-like transporters that span opposing membranes and provide a channel for bulk lipid flow (Kumar *et al*., 2018; Li *et al*., 2020; Dziurdzik and Conibear, 2021; Neuman *et al*., 2022). In many family members, the N-terminus is tethered to the endoplasmic reticulum (ER), while the C-terminus undergoes dynamic recruitment to target membranes, which may regulate organellar lipid transport (Bean *et al*., 2018; Dziurdzik and Conibear, 2021; Leonzino *et al*., 2021).

The subcellular localization of yeast Vps13 varies dynamically according to the state of the cell. Vps13 is present at mitochondria and endosomes during logarithmic growth, but relocates to the nuclear-vacuole junction (NVJ) under starvation conditions or to the prospore membrane during sporulation (Park and Neiman, 2012; Lang *et al*., 2015). This organelle-specific targeting is mediated by distinct membrane adaptors (Bean *et al*., 2018). To date, three such adaptors have been identified – Mcp1, Spo71, and Ypt35 – which direct Vps13 to mitochondrial, prospore, and endosome/vacuolar membranes, respectively (Park *et al*., 2013; John Peter *et al*., 2017; Bean *et al*., 2018).

All three adaptors feature a hydrophobic amino acid-rich “PxP” motif with the consensus sequence Φ-x-x-Φ-x-P-x-P-Φ, where Φ indicates a hydrophobic residue (Bean et al. 2018). These adaptors compete for binding to the Vps13 Adaptor Binding (VAB) domain through their PxP motifs, suggesting they share a common binding site. A recent structural study showed the VAB domain consists of six repeats arranged in an arc, and identified a binding site for the mitochondrial adaptor Mcp1 between the 5^th^ and 6^th^ repeats (Adlakha *et al*., 2022), as predicted from previous studies (John Peter *et al*., 2017; Dziurdzik *et al*., 2020). While other PxP motifs compete with Mcp1 both *in vitro* (Adlakha *et al*., 2022) and *in vivo* (Bean *et al*., 2018; Dziurdzik *et al*., 2020), it remains unclear whether they bind in exactly the same manner. Moreover, none of the three known adaptors can account for Vps13’s role in peroxisome targeting or its participation in the vacuolar protein sorting (VPS) pathway (John Peter *et al*., 2017; Dziurdzik and Conibear, 2021; Yuan *et al*., 2022), suggesting the existence of additional VAB-binding adaptors.

In this study, we demonstrate that the previously defined VAB-PxP motif interface (Adlakha *et al*., 2022) is important for Vps13 targeting *in vivo*. Based on *in silico* modeling and functional analysis, we find that the core PxP motifs from the three known adaptors engage this site through similar contacts. We also identify an extended interface unique to Spo71, suggesting that individual adaptors may fine-tune interactions through additional contacts. Importantly, we discover that the shared VAB-PxP interface is not required for Vps13 function in the VPS pathway, suggesting that Vps13 employs an alternative targeting mechanism to direct vacuolar protein sorting.

## RESULTS

### The three known adaptors share a similar VAB-PxP interface

To identify the Vps13(VAB)-PxP interface for each known adaptor, we used AlphaFold2 (AF2) Multimer-based predictions with ColabFold (Mirdita *et al*., 2022) to model VAB domain repeats 5-6 (Vps13^2321-2543^) together with the PxP motif-containing fragments of each adaptor (Ypt35^1-48^, Mcp1^1-40^, Spo71^381-420^). The resulting high-confidence models, as supported by pLDDT scores and predicted aligned error (PAE) values (Figure S1, A–F), revealed that the 9-residue core PxP motif of each adaptor forms a consistent interface with VAB repeat 6, proximal to repeat 5 (Figure 1, A–D). We next compared these predicted interfaces to the previously resolved crystal structure of the *Chaetomium thermophilum* Vps13(VAB):Mcp1(PxP) complex (Adlakha *et al*., 2022; PDB ID 7U8T) to assess the accuracy of our models.

**Figure 1.**
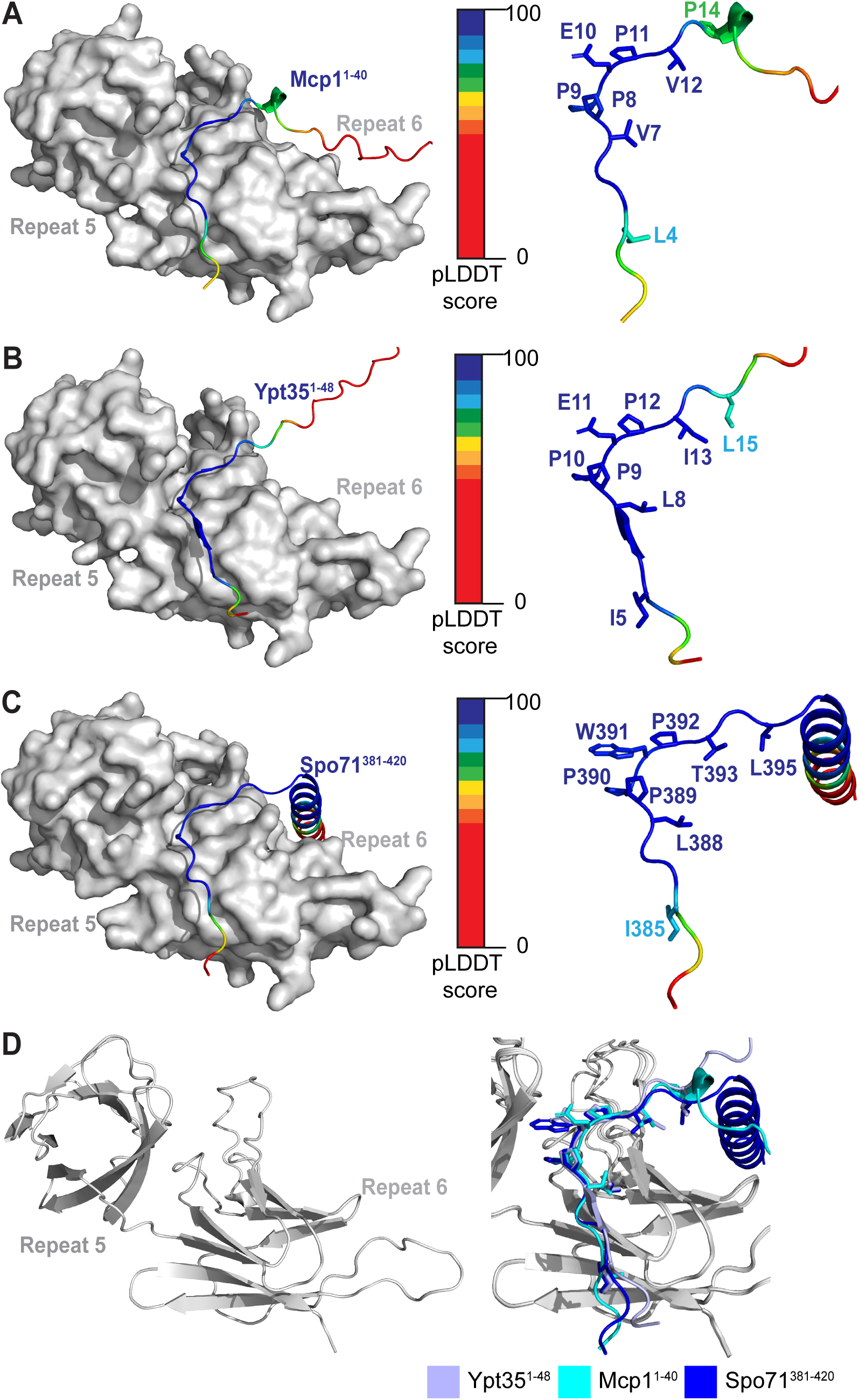
All three adaptors share a predicted interface. (A-C) AlphaFold2-based ColabFold models of VAB domain repeats 5 and 6 interacting with the PxP motifs of (A) Mcp1^1-40^; (B) Ypt35^1-48^; and (C) Spo71^381-420^. PxP motifs are coloured by predicted LDDT score, with VAB domain repeats shown in grey. PxP motif peptides are shown as ribbon diagrams on the right. PxP residues that contact the VAB domain are numbered, and interacting side chains are shown. (D) Aligned ColabFold models show that the predicted interfaces between VAB repeats 5-6 and the PxP motifs of Mcp1, Ypt35, and Spo71 closely match. VAB domain repeats are shown in grey; arbitrary colours are used to distinguish each PxP motif.

To validate our AF2 prediction against the experimentally determined structure, we compared trimmed models corresponding to VAB repeat 5-6 and the 9 core residues of the Mcp1 motif. Both models had favorable geometry based on Molprobity analysis (Chen *et al*., 2010); (Table S1), and displayed strong similarity, with a C⍺ RMSD of 1.196 Å over 157 pruned atoms and similar buried interface areas (613.6 Å^2^ for the predicted model and 591.2 Å^2^ for the experimental structure). The AF2-predicted Vps13(VAB):Ypt35(PxP) and Vps13(VAB):Spo71(PxP) models also had favorable geometry, and comparable interface areas (Table S1). Together, this suggests the predictive models offer a reliable framework for guiding further comparative and functional analyses.

For a more detailed comparison of the predicted and experimental models, we analyzed the buried surface area of the individual interface residues using the PISA server (Krissinel and Henrick, 2007) (Table S2). We found that the consensus residues, which were previously shown to be important for Vps13 membrane recruitment by Ypt35 (Bean *et al*., 2018), make substantial contributions to each interface (Table S2, Figure 1, A-C), and their relative contribution was similar for the different adaptors and yeast species. In general, the same VAB residues interact with the core PxP motifs from all three adaptors, with per-residue buried surface areas comparable to those of the *C. thermophilum* Vps13(VAB):Mcp1(PxP) model. We observed only minor differences: for example, for Mcp1 and Ypt35 – but not Spo71 - the intervening residue between the two conserved prolines (the “x” in “PxP”) was predicted to form a salt bridge with the VAB residue H2397 (Table S2). Collectively, these results demonstrate that the interface between the core PxP motif and the Vps13 VAB domain is largely conserved across different yeast species and different adaptor proteins.

### Residues of VAB domain repeat 6 are critical for maintaining the VAB-PxP interface

Our analysis identified 18 VAB domain residues that interface with all three adaptors: four residues from repeat 5, and fourteen from repeat 6 (Table S2). However, their relative importance for adaptor binding is not known. We previously showed that the second proline (P12) of the PxP motif in Ypt35 is essential for VAB-adaptor interactions (Bean *et al*., 2018). Thus, mutations to VAB residues that interact with this proline are expected to similarly disrupt VAB-PxP binding.

Only four residues were predicted to form an interface with this second proline for all three adaptors: H2397 in repeat 5, and I2464, F2465, and Y2466 in repeat 6 (Table S2). We selected two adjacent residues in VAB repeat 6 (F2465, Y2466) and one residue in repeat 5 (H2397) for mutational analysis (Figure 2, A and B). Mutation of H2397 to alanine had little effect on the Vps13-Ypt35 interaction by co-immunoprecipitation (coIP; Figure 2, C and D). In contrast, the double F^2465^A Y^2466^A (FY^2465-6^A) mutation significantly reduced binding between Ypt35-3HA and Vps13^GFP, which was internally tagged at residue 499 (Lang *et al*., 2015) (24% recovery relative to WT; p<0.0001). This effect was comparable to that observed when mutating both invariant prolines (P^10,12^A) of the Ypt35 PxP motif (19% recovery relative to WT; p<0.001; Figure 2, C and D). These findings support our AlphaFold modeling and confirm the functional importance of the predicted VAB-PxP interface *in vivo*.

**Figure 2.**
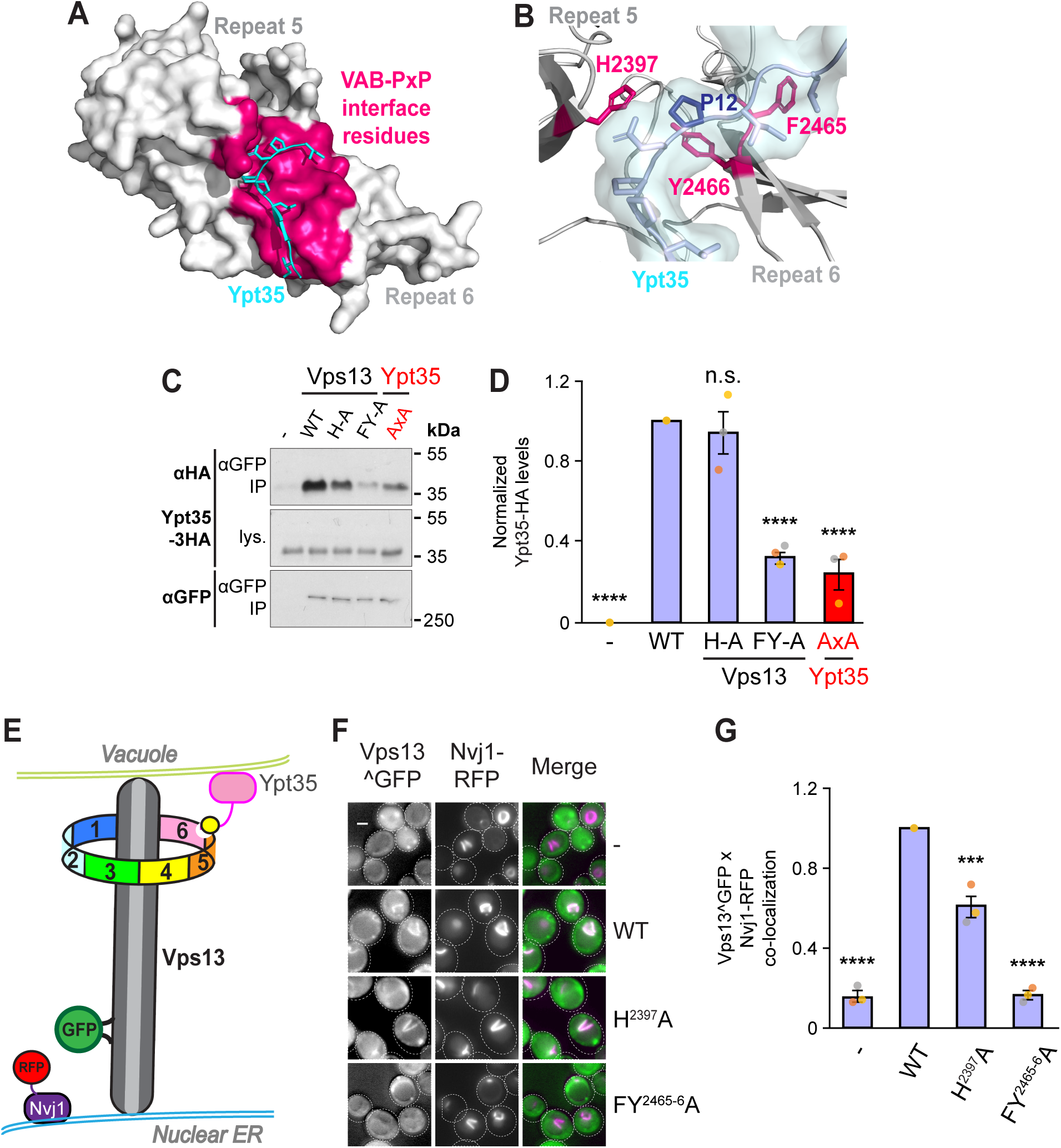
Mutation of the predicted VAB interface disrupts PxP adaptor interactions. (A) AlphaFold2-based ColabFold model of VAB repeats 5 and 6 showing surface residues (hot pink) that are predicted to interact with the core residues of the Ypt35 PxP motif (K4 to I13). (B) Detail of the VAB-PxP interface, highlighting the P12 residue of the Ypt35 PxP motif (navy) and three VAB residues that are predicted to contact it (hot pink). (C-D) Mutation of VAB residues F2465 and Y2466 reduces co-immunoprecipitation of Ypt35-3HA and Vps13^GFP. The co-IP of WT Vps13^GFP and a mutant form of Ypt35 with a P^10,12^A mutation (AxA) is shown as a control (red text). A representative experiment is shown on left (C), with densitometric quantitation of co-purifying HA signal shown on the right (D). All values are relative to WT and are normalized to the levels of HA-tagged protein in the lysate and the amount of Vps13^GFP recovered in the IP. Individual data points are coloured by replicate, n=3. One-way ANOVA test with Dunnett correction, P****<0.0001, n.s. = not significant. Error bars = S.E.M. IP = immunoprecipitate; lys = lysate. “-“ = empty vector, H-A = Vps13 H2397A, FY-A = Vps13 FY^2465-6^A, AxA = Ypt35 P^10,12^A. (E) Schematic of the nuclear-vacuolar junction (NVJ) assay. Under starvation, Ypt35 is recruited to NVJs, which recruits GFP-tagged Vps13. Colocalization at NVJs can be detected by overlap of GFP-tagged Vps13 and RFP-tagged Nvj1. (F) WT Vps13^GFP, but not the FY^2465-6^A mutant, is recruited to the NVJ by Ypt35-3HA after growth in acetate media. The RFP channel shows the localization of the NVJ marker Nvj1-RFP. Bar, 2µm. (G) Automated quantitation shows the fraction of cells containing NVJ-localized Vps13^GFP relative to WT, n=3, ≥616 cells/strain/sample. Individual data points are coloured by replicate. One-way ANOVA test with Dunnett correction, P***<0.001, P****<0.0001. Error bars = S.E.M.

We next examined whether these VAB residues are required for the recruitment of endogenous Vps13 to the nuclear-vacuolar junction (NVJ), a membrane contact site between the nuclear ER and vacuole. Vps13 is recruited to NVJs by Ypt35 under starvation conditions (Lang *et al*., 2015; Bean *et al*., 2018). Using automated quantification, we assessed the co-localization between mutant forms of Vps13^GFP and the NVJ marker Nvj1-RFP (Lang *et al*., 2015; Bean *et al*., 2018) in cells overexpressing Ypt35-3HA (Figure 2, E-G). Both H^2397^A and FY^2465-6^A mutations significantly reduced the NVJ localization of Vps13 relative to WT (p<0.0001). The effect of the FY^2465-6^A mutation was comparable to mutating the invariant prolines (P^10,12^A) in Ypt35, and substantially stronger than the H^2397^A mutation. These results demonstrate that the VAB domain residues in the predicted PxP motif-binding interface are critical for Vps13 localization to membrane contact sites *in vivo*.

To determine whether these VAB domain residues are necessary for Vps13^GFP membrane recruitment by each of the three adaptors, we used a chimeric endosome recruitment assay (Bean et al., 2018). In this assay, each PxP peptide was fused to a PI3P-binding, endosome-targeted RFP-FYVE module in a strain lacking the endogenous endosomal adaptor, Ypt35 (Figure 3A). By quantifying the overlap of GFP and RFP puncta (Figure 3, B-G), we determined that the FY^2465-6^A mutations significantly inhibit the endosomal recruitment of Vps13^GFP by each PxP motif to <25% of WT levels. Together, these results demonstrate that mutation of the predicted interface in VAB domain repeat 6 effectively blocks adaptor-mediated Vps13 recruitment to membranes.

**Figure 3.**
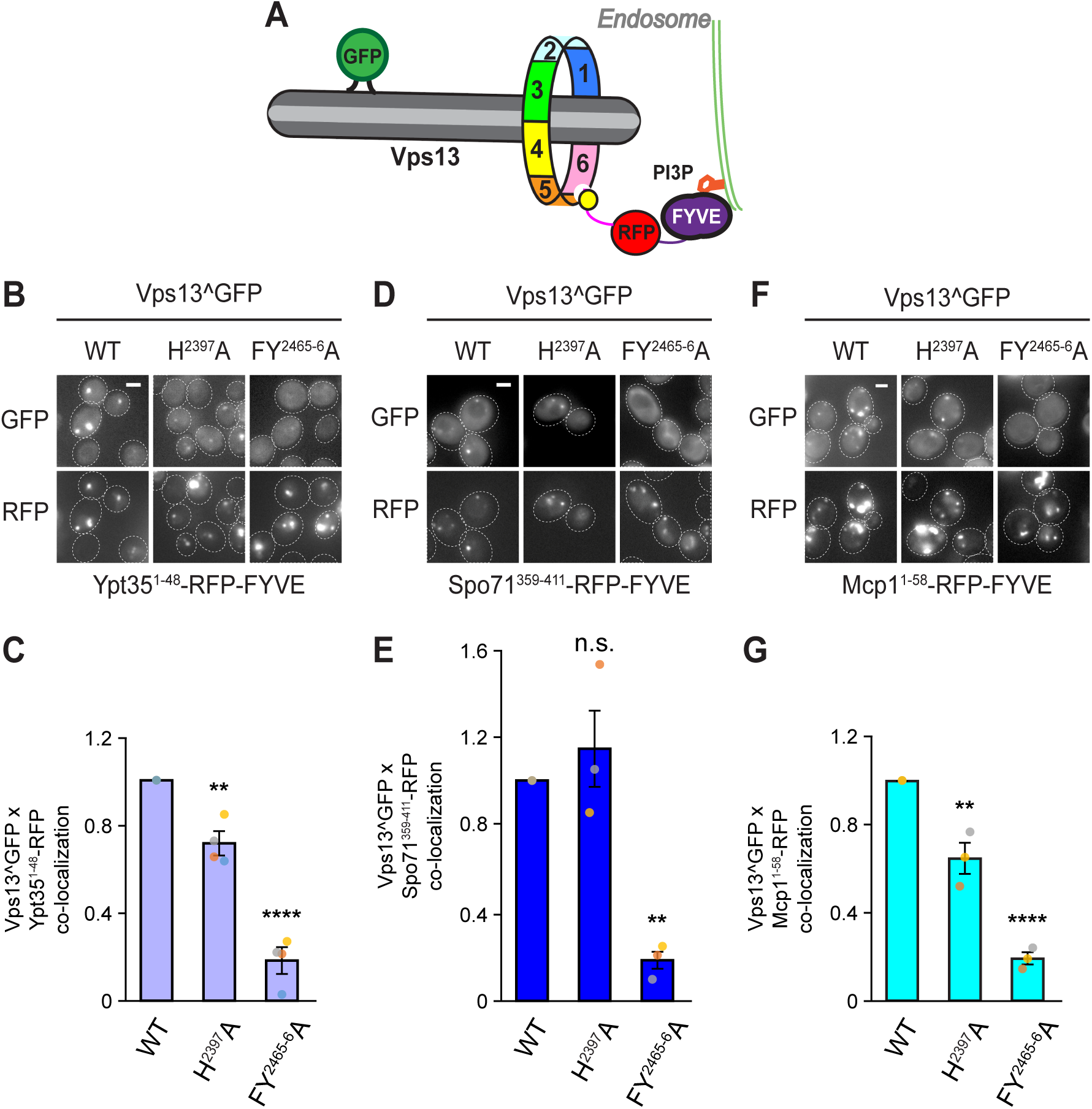
VAB mutations disrupt recruitment by PxP adaptors. (A) Schematic of PxP motif chimeric recruitment assay. The PxP motif (yellow dot) is attached to an RFP-tagged FYVE domain, which localizes to the endosome. Vps13^GFP is recruited to the endosome by the motif, and colocalization can be evaluated from overlap of GFP and RFP puncta. (B, D, and F). Co-localization microscopy of plasmid-expressed WT and mutant forms of Vps13^GFP co-expressed with chimeric molecules consisting of the endosome-targeted RFP-FYVE domain fused to adaptor PxP motifs: (B) Ypt35^1-48^, (D) Spo71^359-411^, and (F) Mcp1^1-58^. All plasmids were expressed in *ypt35*Δ yeast cells. Bar, 2µm. (C, E, and G) Automated quantitation of co-localizing GFP and RFP puncta, relative to total number of RFP puncta per sample. Values are normalized to WT. n⩾3; ⩾208 cells/strain/replicate. One-way ANOVA test with Dunnett correction, P**<0.01, P****<0.0001, n.s. = not significant. Error bars = S.E.M.

The H^2397^A mutation modestly inhibited the recruitment of Vps13^GFP by the PxP motifs from Ypt35 and Mcp1, but not Spo71 (Figure 3, B-G). This is consistent with the predicted involvement of this residue in specifically recognizing the PxP motifs of these two adaptors, and suggests the recruitment assay may be more sensitive than co-immunoprecipitation for detecting subtle differences in Vps13-adaptor interactions. We used this sensitive chimeric recruitment assay to determine if the effect of the H^2397^A mutation is enhanced by point mutations in the Ypt35 PxP motif (Bean *et al*., 2018), but found no conclusive evidence of synergistic defects (Figure S2, A and B).

Another VAB domain repeat 5 residue, R2396, which was suggested to contribute to the VAB interface with Mcp1 (Adlakha *et al*., 2022), contacted the PxP motif of only Ypt35 and Mcp1 in our models. Using the chimeric recruitment assay, we found no effect of the R^2396^A mutation, either alone or in combination with H^2397^A (Figure S3, A and B). Taken together, this suggests that the VAB domain repeat 5 residues do not make critical contributions to the VAB-PxP interface.

### Spo71 has a unique extended interface with the VAB domain

The known PxP adaptors compete with each other for Vps13 binding (Bean *et al*., 2018), but how this competition is regulated is not well understood. Possible mechanisms include post-translational modifications of Vps13 or its adaptors, changes in relative expression levels, or subtle differences in the VAB-PxP interface that confer stronger binding interactions.

While analyzing the interface between VAB repeat 6 and the Spo71 PxP motif, we found that an adjacent alpha helix in Spo71 is also predicted to interact with VAB repeat 6 (Figure 4A). Analysis of the model interfaces using PISA (Krissinel and Henrick, 2007) identified four Spo71 residues and six Vps13 residues that are present at this second interface (Table S3). Although one of these Vps13 residues, I2435, is predicted to interact with the core PxP motif of Ypt35, but not that of Spo71, the remaining Vps13 residues are predicted to interact specifically with the Spo71 helix (Figure 4B).

**Figure 4.**
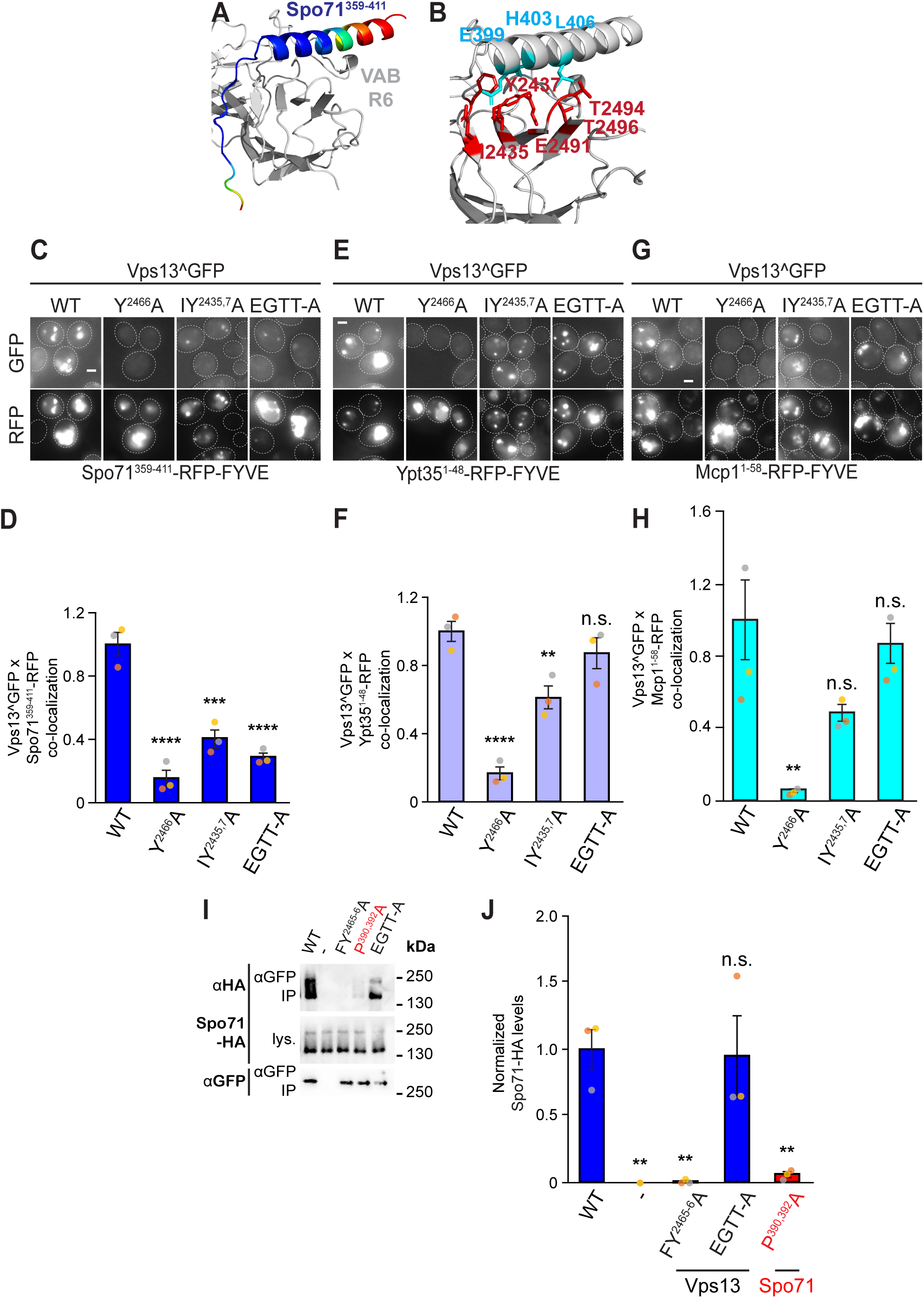
The VAB domain has a unique partial interface with Spo71. (A) AlphaFold2-based ColabFold model of VAB repeats 5 and 6 interacting with Spo71381-420, highlighting the alpha helix C-terminal to the PxP motif. Spo71 coloured by pLDDT score. (B) Residues of the interface predicted to interact (light blue = Spo71 residues, red = Vps13 residues). (C, E, and G) Co-localization microscopy of plasmid-expressed WT and mutant forms of Vps13^GFP co-expressed with chimeric molecules of the endosomally-targeted RFP-FYVE domain fused to adaptor PxP motifs: (C) Spo71^359-411^, (E) Ypt35^1-48^, and (G) Mcp1^1-58^. All plasmids were expressed in *ypt35*Δ cells. Bar, 2µm. (D, F, and H) Automated quantitation of co-localizing GFP and RFP puncta, relative to total number of RFP puncta per sample. Values are normalized to WT. n≥3; ≥171 cells/strain/replicate. One-way ANOVA test with Dunnett correction, P**<0.01, P***<0.001, P****<0.0001, n.s. = not significant. (I) Mutation of VAB residues F2465 and Y2466 reduces co-immunoprecipitation of Spo71-HA and Vps13^GFP, while the Vps13 E^2491^A GT^2493-4^A T^2496^A VAB mutation that targets the Spo71 alpha helix interface does not significantly reduce binding. The Spo71 P^390,392^A mutation that disrupts the Spo71 PxP motif is shown as a control (red text). “-“ indicates an empty vector control. (J) Quantitation of coIP data. All values are relative to WT coIP levels. Individual data points are coloured by replicate, n=3. One-way ANOVA test with Tukey correction, P**<0.01, n.s. = not significant. IP = immunoprecipitate; lys = lysate. EGTT-A = E^2491^A GT^2493-4^A T^2496^A. Error bars = S.E.M.

To determine if mutations at the VAB repeat 6-Spo71 helix interface affect Vps13 recruitment, we used our chimeric recruitment assay (see Figure 3). We introduced two sets of mutations in Vps13: one set targeting residues predicted to contact only the Spo71 alpha helix (E^2491^A, GT^2493-4^A, and T^2496^A, referred to as EGTT-A) and another set targeting residues that could affect binding to other adaptors (I^2435^A and Y^2437^A). We tested both sets of mutations with all three adaptor motifs to determine if the interfaces are adaptor-specific.

The I^2435^A Y^2437^A mutations, which are also predicted to impact binding of the Ypt35 core PxP motif, significantly decreased recruitment of Vps13 by both Spo71 and Ypt35. In contrast, the EGTT-A mutations specifically reduced recruitment by the Spo71 PxP motif (Figure 4, C-H). This finding suggests the predicted VAB-Spo71 helix interface contributes to the recognition of Vps13 by Spo71.

We further assessed the importance of the Spo71 alpha helix-Vps13 interface by co-immunoprecipitation. As expected, mutations disrupting the VAB-PxP interface (FY^2465-6^A) significantly reduced binding, as did mutation of the Spo71 invariant prolines (P^390,392^A). However, the EGTT-A mutation did not cause a significant change in binding (Figure 4, I and J). This discrepancy suggests that disruption of the VAB-Spo71 helix interface has a relatively modest effect that can only be detected by the more sensitive colocalization-based recruitment assay.

### Predicted Vps13(VAB)-PxP binding residues are not required for CPY sorting

None of the adaptors identified to date are required for Vps13’s function in the VPS pathway, which sorts newly synthesized vacuolar proteins such as carboxypeptidase Y (CPY) to the vacuole (Conibear and Stevens, 1998; Dziurdzik *et al*., 2020; Dziurdzik and Conibear, 2021; Figure 5A). To test if the PxP motif binding interface is important for CPY sorting, we expressed Vps13^GFP carrying mutations that disrupt binding to either the PxP motif (H^2397^A or FY^2465-6^A), and compared these to previously described VAB mutations (Dziurdzik *et al*., 2020) that alter invariant asparagines in repeat 1 (N^1875^A, referred to as N1A) or repeat 6 (N^2428^A, referred to as N6A) (Figure 5, B and C). CPY was secreted from the Vps13 N1A or N6A mutants, as previously reported (Dziurdzik *et al*., 2020). However, CPY secretion from the H^2397^A single mutation and FY^2465-6^A double mutation was near background levels (Figure 5, B and C). This suggests that the putative Vps13 adaptor that functions in the VPS pathway is unlikely to use a PxP motif, but instead may interact with the VAB domain through a distinct interface.

**Figure 5.**
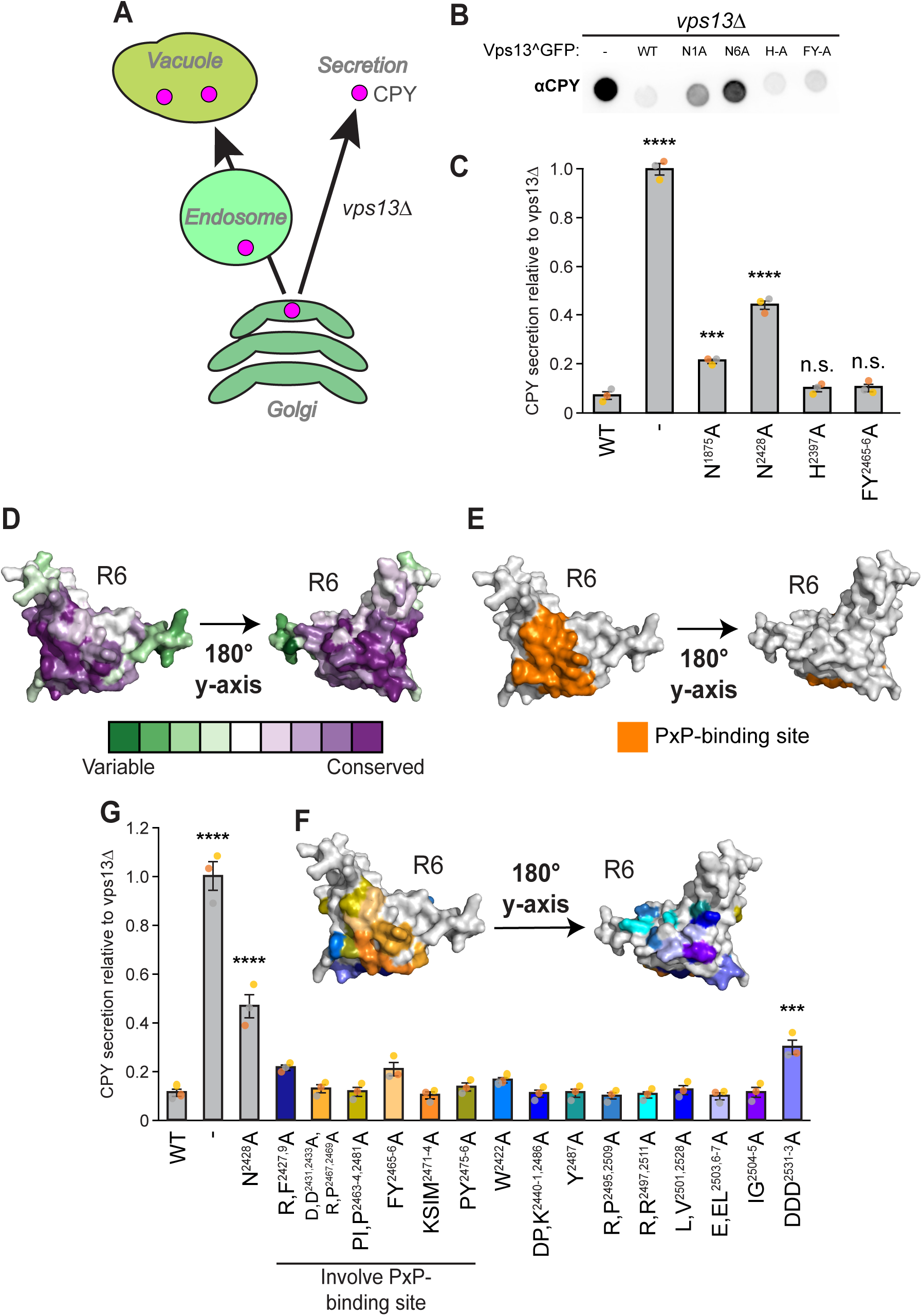
VAB-PxP interface does not impact CPY secretion. (A) Schematic of the CPY sorting pathway. In wild type cells, newly synthesized carboxypeptidase Y (CPY) is shuttled from the Golgi to the vacuole via the endosome. Upon deletion of VPS13, CPY is instead secreted from the cell, where it can be detected by Western blot when cells are grown on nitrocellulose membranes. (B) Spot assay for CPY secretion, showing effects of mutating Vps13 invariant asparagines in repeats 1 (N1: N1875) and 6 (N6: N2428), or VAB-PxP interface residues (H-A: H2397A and FY-A: FY2465-6A). WT and mutant forms of Vps13^GFP were expressed from plasmids in *vps13*Δ mutant cells; “-“ = empty vector control. (C) Deletion of VPS13 or mutation of the invariant asparagines causes carboxypeptidase Y secretion, in contrast to the VAB-PxP interface mutations H^2397^A and FY^2465-6^A. WT or mutant forms of Vps13^GFP were expressed from plasmids in *vps13*Δ mutant cells; CPY secretion was quantified from strains grown as 1536-colony arrays. All values are normalized to a *vps13*Δ control. Individual data points are coloured by replicate, n=3; ≥24 technical replicates per strain. One-way ANOVA test with Tukey correction, P***<0.001. P****<0.0001, n.s. = not significant. (D, E, and F) AlphaFold2 models of VAB repeat 6 (Vps13^2421-2543^) with residues highlighted to show: (D) conservation scores based on ConSurf, (E) residues present at the PxP motif interface, (F) residues that were mutated to alanine and tested for CPY secretion. Mutated residues at the PxP motif interface are shown in orange/brown, while residues outside this interface are shown in shades of blue/purple (G) Quantitation of CPY secretion from vps13Δ strains expressing wild type and mutant forms of Vps13 and grown as 1536-colony arrays. All values are normalized to a *vps13*Δ control. Individual data points are coloured by biological replicate, n=3; ≥24 technical replicates per strain. One-way ANOVA test with Tukey correction, P***<0.001, P****<0.0001, only significant differences are shown. Displayed stats are relative to WT. Error bars = S.E.M.

### Scanning mutagenesis of VAB repeat 6 does not identify a site exclusive to the CPY sorting pathway

Two previous observations indicated that a hypothetical adaptor for the VPS pathway might bind to a site on VAB repeat 6. First, a higher level of CPY missorting is observed by mutation of the invariant asparagine of repeat 6 relative to that of repeat 1 (Figure 5, B and C) (Dziurdzik *et al*., 2020). Second, expression of high levels of soluble PxP peptides fused to mCherry also causes some CPY missorting (Dziurdzik *et al*., 2020), suggesting this competitively inhibits the binding of the putative VPS adaptor (Figure 5A). Thus, we hypothesized that the VPS adaptor binds at a site that is close to or partially overlapping the PxP motif binding site.

Using ConSurf analysis (Ashkenazy *et al*., 2016), we mapped highly conserved surface residues of VAB repeat 6 (Figure 5D). This revealed several conserved surface patches not involved in binding to the core PxP motif (Figure 5E). We systematically mutated these conserved surface patches, focusing on residues proximal to each other in the predicted 3D structure (Figure 5F). Using the same CPY overlay assay (Figure 5C), we found that mutation of one specific patch, which changed three adjacent aspartic acids to alanines (DDD^2531-3^A), significantly increased CPY secretion above wild-type levels (2.5-fold, p<0.001) (Figure 5G).

To further define the critical region, we generated and tested additional mutations, including mutations of surrounding residues (e.g. DDDMQ^2531-5^A) and charge-swap mutations that converted the aspartic acids to lysines (DDD^2531-3^K) (Figure 6. A and B). The DDD^2531-3^K mutation exhibited the strongest effect of any Vps13 missense mutation tested (roughly 40% of *vps13* null, p<0.0001). Western blot analysis confirmed that these mutant Vps13 proteins were expressed at levels comparable to wild type (Figure 6, C and D).

**Figure 6.**
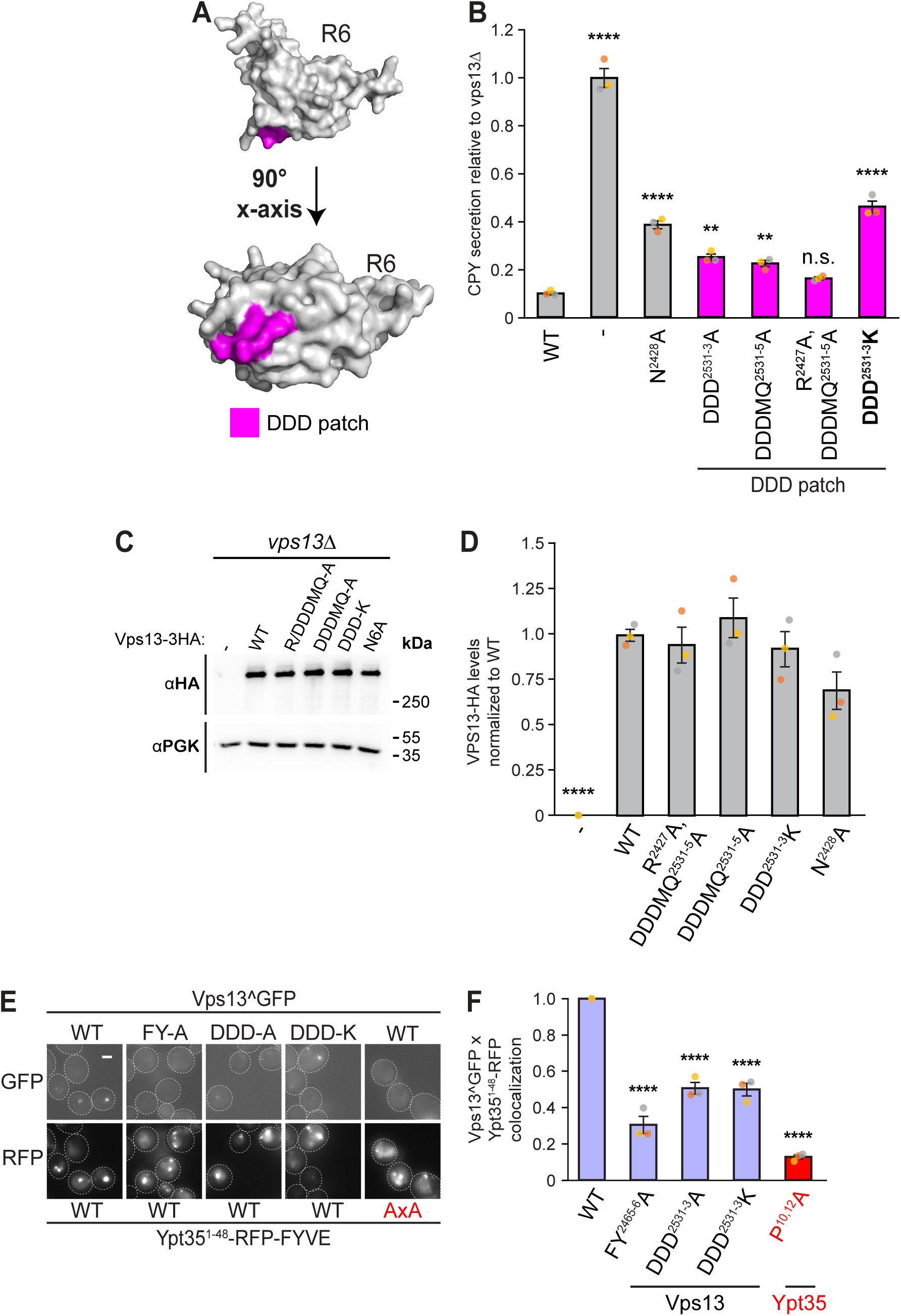
A Vps13 DDD patch contributes to CPY sorting and PxP-based membrane recruitment. (A) AlphaFold2-based ColabFold of Vps13 VAB R6 (Vps13^2421-2543^) with residues DDD2531-3, which form a surface patch, highlighted in pink. (B) Quantitation of CPY secretion from *vps13*Δ strains expressing wild type and mutant forms of Vps13 with a C-terminal 3HA tag and grown as 1536-colony arrays. All values are normalized to a *vps13*Δ control. Individual data points are coloured by biological replicate, n=3; n≥24 technical replicates per sample. One-way ANOVA test with Tukey correction, P***<0.001, P****<0.0001, only significant differences are shown. Displayed stats are relative to WT. “-“ = empty vector control. (C) Effects of mutating DDD patch residues on Vps13-3HA stability. A representative experiment is shown on left (C), with densitometric quantitation of HA signal shown on the right (D), n=3. One-way ANOVA test with Dunnett correction, P****<0.0001. Only significant values are shown. All values relative to WT. (E) Co-localization microscopy of plasmid-expressed WT and mutant forms of Vps13^GFP co-expressed with chimeric molecules consisting of the endosomally-targeted RFP-FYVE domain fused to the PxP motif of Ypt35. Bar, 2µm. (F) Automated quantitation of co-localizing GFP and RFP puncta, relative to total number of RFP puncta per sample. Values are normalized to WT, n≥3; ≥87 cells/strain/replicate. One-way ANOVA test with Dunnett correction, P****<0.0001. Error bars = S.E.M.

To determine whether the effect of these mutations was specific to CPY secretion, we examined their impact on Vps13 recruitment to endosomes by the Ypt35 PxP motif-containing chimeric protein. Both the DDD^2531-3^A and DDD^2531-3^K mutations significantly reduced overlap between Vps13^GFP and Ypt35^1-48^-RFP-FYVE puncta (to 51% and 50% of WT levels, p<0.0001, respectively) (Figure 6, E and F). Because these mutations also impact PxP adaptor-mediated localization, yet lie outside the VAB:PxP interface, this cluster of aspartic acid residues likely plays a broader role in Vps13 conformation or function, rather than a role specific to vacuole protein sorting. Overall, the fact that our comprehensive mutational scanning of repeat 6 did not identify a VPS-specific site suggests that the putative VPS adaptor does not simply use a divergent motif that binds at a nearby or overlapping site. Instead, our results suggest the targeting of Vps13 to organelles relevant for the VPS pathway uses a distinct targeting mechanism.

## DISCUSSION

Using AlphaFold-based protein structure prediction, we modeled interactions between the yeast Vps13 VAB domain and the PxP targeting motifs from three different organelle adaptors. Mutational analysis confirmed that the predicted VAB-PxP interface shared by all three adaptors is functionally important for Vps13 localization *in vivo*. Unexpectedly, we found that this shared interface is not required for Vps13’s role in the vacuolar protein sorting pathway, suggesting the existence of alternative, previously unrecognized targeting mechanisms.

Our predictive modeling strongly supports the idea of a single, shared binding site for PxP adaptor motifs that involves just two of the six VAB repeats, consistent with previous biochemical and structural studies (John Peter *et al*., 2017; Bean *et al*., 2018; Adlakha *et al*., 2022). Indeed, our modeling suggests that the consensus PxP motif residues from Mcp1, Ypt35, and Spo71 all engage this interface in a similar manner, and we found that altering just two VAB domain residues predicted to contact the critical second proline of the PxP motif effectively disrupted interactions with all three adaptors. These results highlight both the conserved nature of the PxP binding mechanism and the central importance of the proline residues in mediating adaptor recognition.

While the core binding sites proved remarkably similar, we identified an alpha helix in Spo71 located C-terminal to the core PxP motif that forms additional contacts with VAB repeat 6. Previous studies had suggested that Spo71 exhibits stronger binding than other adaptors (Bean *et al*., 2018), and this extended interface could theoretically enhance binding and provide Spo71 with a competitive advantage. Although mutations at this extended interface had only modest effects on Spo71-Vps13 binding, the finding demonstrates that PxP adaptors can use overlapping yet distinct binding strategies to achieve specificity.

The discovery that disrupting the shared PxP binding site does not impair Vps13’s role in vacuolar protein sorting was unexpected. Given the importance of organelle-specific adaptors for Vps13 targeting, we had expected that eliminating PxP binding would broadly impair Vps13 function. Instead, our results reveal a clear functional separation: while PxP adaptors regulate Vps13’s targeting to mitochondria, prospore membranes, and endosomes, the VPS pathway appears to rely on entirely different targeting mechanisms.

We initially hypothesized that the VPS pathway uses a divergent adaptor that lacks the characteristic proline residues but still binds at a site on repeat 6 close to, or overlapping, the core PxP interface. To test this possibility, we conducted extensive mutational scanning of conserved VAB residues outside the established PxP binding site. Rather than identifying a VPS-specific binding patch, this analysis revealed residues whose mutation impaired both CPY sorting and PxP-mediated localization. This suggests that these mutations cause broader changes that affect multiple interaction sites.

Based on these observations, we propose that the putative VPS adaptor binds at a distinct site that is influenced by the conformation of the VAB domain. In this model, either occupying the PxP site or disrupting the folding of repeat 6 can change the properties of the VAB domain, indirectly affecting VPS adaptor binding. A related idea was proposed by the Neiman laboratory (Park *et al*., 2021), who found that mutation of Ypt35 and the conserved asparagine residue in repeat 6 synergistically contribute to CPY missorting, and suggested that conformational changes in repeat 6 impact the binding of a co-adaptor that functions redundantly with Ypt35. Several domains within Vps13 have also been shown to bind specific lipids and contribute to membrane targeting (De *et al*., 2017; Kolakowski *et al*., 2020; Hanna *et al*., 2023). This raises the possibility that lipid binding works in conjunction with protein adaptors to achieve proper localization.

The role of Vps13 in the VPS pathway suggests that it supplies lipids to a key organelle in this this pathway to support membrane remodeling events, but the identity of this organelle is not known. Recent studies have linked Vps13 to the ESCRT machinery, which generates intralumenal vesicles during multivesicular body formation at endosomes (Henne *et al*., 2011). Efficient intralumenal vesicle formation requires Vps13 recruitment to a membrane contact site between the endosome and the ER and/or lipid droplets (Suzuki *et al*., 2024; Gao *et al*., 2025). At these sites, Vps13 may supply lipids to the endosome membrane, where it partners with the Any1 scramblase to ensure lipid equilibration across membrane leaflets needed for efficient ILV formation (Gao *et al*., 2025).

Is this the key role of Vps13 in the VPS pathway? Loss of Ypt35 causes the same aberrant endosomal morphology as loss of Vps13 (Bean *et al*., 2018; Gao *et al*., 2025), yet Ypt35 is dispensable for CPY sorting. This suggests that Vps13 acts at a distinct, Ypt35-independent, trafficking step. Vps13 has been implicated in the homotypic fusion of late Golgi and the fusion of transport vesicles with the late endosome using a cell-free assay (De *et al*., 2017), and is important for early endosomal recycling (Dalton *et al*., 2017), suggesting that Vps13 has additional roles beyond ILV formation.

The importance of the VAB domain in binding targeting adaptors appears to be conserved in humans (Hanna *et al*., 2023; Tornero-Écija *et al*., 2023; Du *et al*., 2024; Wang *et al*., 2025). VAB-binding adaptor proteins have been identified for VPS13A, VPS13B and VPS13C (Tornero-Écija *et al*., 2023; Du *et al*., 2024; Wang *et al*., 2025) though it is not known if these proteins use the Φ-x-x-Φ-x-P-x-P-Φ consensus motif found in yeast adaptor proteins. Human VPS13 proteins can also bind to targeting adaptors and lipid scramblases at sites outside the VAB domain (Park and Neiman, 2020; Guillen-Samander *et al*., 2021; Dall’Armellina *et al*., 2023; Hanna *et al*., 2023; Covill-Cooke *et al*., 2024). Intriguingly, there is evidence that even in these cases, the VAB domain contributes to targeting (Park and Neiman, 2020; Guillen-Samander *et al*., 2021; Park *et al*., 2022). The recent demonstration in human cells that VAB binding to Rab7 is subject to autoinhibition further supports the concept that conformational changes within the VAB domain are important for regulating VPS13 targeting and function (Wang *et al*., 2025). Further studies are needed to understand how Vps13 is targeted to, and functions at, diverse organelle membranes and how the VAB domain contributes to the intracellular localization of VPS13 proteins in eukaryotes.

## MATERIALS AND METHODS

### Yeast strains and plasmids

Strains, plasmids and primers used in this study are described in Supplementary Tables S4-S6. All strains from this study were made by polymerase chain reaction (PCR)-based homologous recombination (Longtine *et al*., 1998; Janke *et al*., 2004). Gene deletions were confirmed by colony PCR. Plasmids expressing mutant versions of Vps13^GFP (pKJ068-pKJ077 and pKJ118) and VAB-ENVY (pMD304 and pMD323) were made by homologous recombination in yeast by co-transforming linearized plasmids with PCR products containing the desired mutations, generated by primer extension. To generate mutations in Vps13 repeat 6, sequences encoding residues 2414-2541 were first replaced by a PCR-generated superfold-GFP stuffer using homologous recombination with a SalI-digested pRS416-VPS13-3HA vector (pMD549) to create pMD550. pRS416-VPS13-3HA derivatives were then made by digesting pMD550 with XbaI and SalI and replacing the sfGFP stuffer with a synthetic wildtype or mutated VPS13(aa2414-2541) sequences (TWIST Bioscience, Wilsonville, OR). All plasmids were recovered in *Escherichia coli* and sequenced.

### AlphaFold predictions and analysis

Prediction of protein structure and binding interfaces was performed using the AlphaFold2-powered ColabFoldv2 (Jumper *et al*., 2021; Mirdita *et al*., 2022) VAB domain repeats 5-6 models were generated using amino acid boundaries of 2321-2543. PDBePISA (Krissinel and Henrick, 2007) was used to identify interfaces. Conservation analysis was conducted using an AlphaFold2-predicted model of Vps13 (aa1251-3144) submitted to the ConSurf server (Ashkenazy *et al*., 2016) with default settings. PyMOLv2.5.4 was used to generate models that were assembled in Adobe Illustrator v27.2.

### PISA and MolProbity analyses to identify interfaces and model geometry

PyMOLv2.5.4 was used to extract structures corresponding to VAB domain repeats 5-6 and the PxP motifs from either the *C. thermophilum* VAB-Mcp1 crystal structure (PDB 7U8T; *C. thermophilum* Vps13^2413-2626^ and Mcp1^17-25^) or the *S. cerevisiae* AlphaFold amber-relaxed VAB-PxP models (Vps13^2321-2543^ with Ypt35^5-13^, Mcp1^4-12^, Spo71^385-393^ or Spo71^381-420^). These models were analyzed by Molprobity (Chen *et al*., 2010) and PDBePISA (Krissinel and Henrick, 2007) using the default settings. Comparisons between *C. thermophilum* and *S. cerevisiae* VAB domain sequences were conducted using the Jalview multiple sequence alignment tool (Waterhouse *et al*., 2009).

### Co-immunoprecipitation and stability assays

For co-immunoprecipitation experiments, yeast were grown to log phase in minimal synthetic dextrose media and 70 OD_600_ of cells stored at −80°C until further use. Cells were thawed and resuspended in 500 μl lysis buffer (50 mM HEPES pH 7.4, 0.1% Tween-20, 50 mM NaCl, 1 mM EDTA, 1 mM PMSF, and 1× yeast/fungal ProteaseArrest (G-Biosciences) and lysed by vortexing with 100µl 0.5 mm diameter glass beads. 50 μl of lysate was mixed with 2x Laemmli Sample Buffer (240 mm Tris-Cl pH 6.8, 8% SDS, 40% glycerol, 0.004% bromophenol blue, 10% β-mercaptoethanol) containing 8 M urea. The remaining 450 µl lysate was incubated with polyclonal rabbit anti-GFP (EU2; Eusera) for 1 h followed by Protein A-Sepharose beads (GE Healthcare) for one hour at 4°C. Washed beads were resuspended in 50 μl of Thorner buffer (40 mm Tris pH 6.8, 8 M urea, 5% SDS, 0.1 M EDTA, 1% β-mercaptoethanol, 0.4 mg/ml bromophenol blue) and proteins eluted by heating at 100°C for 5 min. Samples were run on SDS-PAGE gels, transferred to nitrocellulose membranes, and blotted with monoclonal mouse anti-GFP (11814460001; Roche) or monoclonal mouse anti-HA (MMS-101R; Covance), and horseradish peroxidase-conjugated polyclonal goat anti-mouse secondary antibody (115–035-146; Jackson ImmunoResearch Laboratories).

Blots were developed with Amersham ECL (GE Healthcare) or Amersham ECL Prime (GE Healthcare) reagents and exposed to Amersham Hyperfilm (GE Healthcare) which was scanned. Alternatively, chemiluminescence was detected with the Vilber Fusion FX (Marne-la-Vallee, France). Densitometry was conducted in ImageJ (Schneider *et al*., 2012). Significance was determined by one-way ANOVA followed by Tukey multiple comparison test, assuming normal distribution, using GraphPad Prism 10.1.1.

For stability analysis, 50µl cell lysate, generated as above, was mixed with 50µl 2x Laemmli Sample Buffer, run on SDS-PAGE gels, and subjected to western blotting as described above.

### Fluorescence microscopy and automated quantitation

Fluorescence microscopy was conducted as previously described (Bean *et al*., 2018). Yeast were grown at room temperature in synthetic dextrose-based media to log phase, and transferred to 0.1mg/ml concanavalin A–treated glass bottom MatriPlates (Brooks). To localize proteins to NVJs, 2 OD/ml of log phase yeast were grown for 18 h at 30°C in synthetic complete media with 2% acetate, and 150µl fresh synthetic complete with acetate media was added to the MatriPlates (Brooks Life Science Systems) prior to imaging. Imaging was conducted at room temperature on a DMi8 microscope with a high-contrast Plan Apochromat 63×/1.30 Glyc CORR CS objective (Leica Microsystems GmbH), an ORCA-Flash4.0 digital camera (Hamamatsu Photonics), and MetaMorph 7.8 software (MDS Analytical Technologies). Measurements were taken on raw images. Images were adjusted for figure generation with MetaMorph, and image cropping was performed in Adobe Photoshop v23.2 and Adobe Illustrator v27.2.

Cellular puncta and NVJs were analyzed using custom MetaMorph 7.8 journals. Live and dead cells were identified through set thresholds using the Count Nuclei function. Dead cells and other non-cellular fluorescent signals were masked using the LogicalAND function. The number of puncta was determined with the Granularity function. To identify NVJs, bright objects were located with TopHat (Nvj1-RFP) while TopHat and Integrated Morphometry Analysis was used to identify weaker signals (Vps13^GFP) based on size and shape. To measure colocalization, a LogicalAND function was used to detect if previously defined objects in GFP and RFP channels occupied the same space, after masks were applied. For non-NVJ colocalization assays, colocalization was normalized to total number of RFP puncta. Significance was determined by one-way ANOVA followed by Dunnett multiple comparison test or two-way ANOVA Tukey multiple comparison test (Figure S2B), assuming normal distribution, using GraphPad Prism 10.1.1.

### CPY secretion

Secreted CPY was captured on nitrocellulose membranes using two different approaches. For array-based measurements, *vps13* null yeast expressing 3HA-tagged Carboxypeptidase Y from the endogenous *PRC1* locus were transformed with wildtype or mutant forms of pVPS13^GFP or pRS416-VPS13-3HA. Strains were grown on selective media in 96-colony format on rectangular plates, then subsequently pinned to 384, then 1536 format using a BM3-BC pinning robot (S&P Robotics inc.). Colonies were transferred to nitrocellulose that was placed on a selective media plate and grown overnight at 30°C. Spot-based CPY secretion assays were conducted as previously described (Dziurdzik *et al*., 2020). Briefly, 1 OD_600_/ml of each log-phase yeast culture was serially diluted and spotted on minimal synthetic dextrose media plates. The plates were covered with nitrocellulose and incubated overnight at 30°C.

Nitrocellulose membranes were washed to remove cells, blocked with 5% skim milk in 1x TPBS (0.1% Tween-20, 137mM NaCl, 2.7mM KCl, 10mM Na_2_HPO_4_, 1.8mM KH_2_PO_4_, pH 7.4), and incubated with monoclonal mouse anti-HA (MMS-101R Covance) (array assays) or monoclonal mouse anti-carboxypeptidase Y (A-6428; Thermo Fisher Scientific) (spot assays), followed by horseradish peroxidase-conjugated polyclonal goat anti-mouse (115–035-146; Jackson ImmunoResearch Laboratories). Signal was detected after incubation with Amersham ECL (GE Healthcare) reagents and imaged with the Vilber Fusion FX. Resulting images were cropped and adjusted with Adobe Photoshop v23.2 and Adobe Illustrator v27.2. Intensity of imaged colonies was analyzed from array-based assays using CellProfiler (Lamprecht *et al*., 2007). Significance was determined by one-way ANOVA followed by Tukey multiple comparison test, assuming normal distribution, using GraphPad Prism 10.1.1.

## ACKNOWLEDGEMENTS

This work was supported by funding from the Canadian Institutes of Health Research (grants 177941 and 180544 to EC, and Frederick Banting and Charles Best Canada Graduate Doctoral Scholarships 170920 to SKD and 181571 to KRJ).

## Non-standard abbreviations

VAB: Vps13 adaptor binding
NVJ: nucleus-vacuole junction

## SUPPLEMENTARY MATERIAL

**Supplemental Figure 1.**
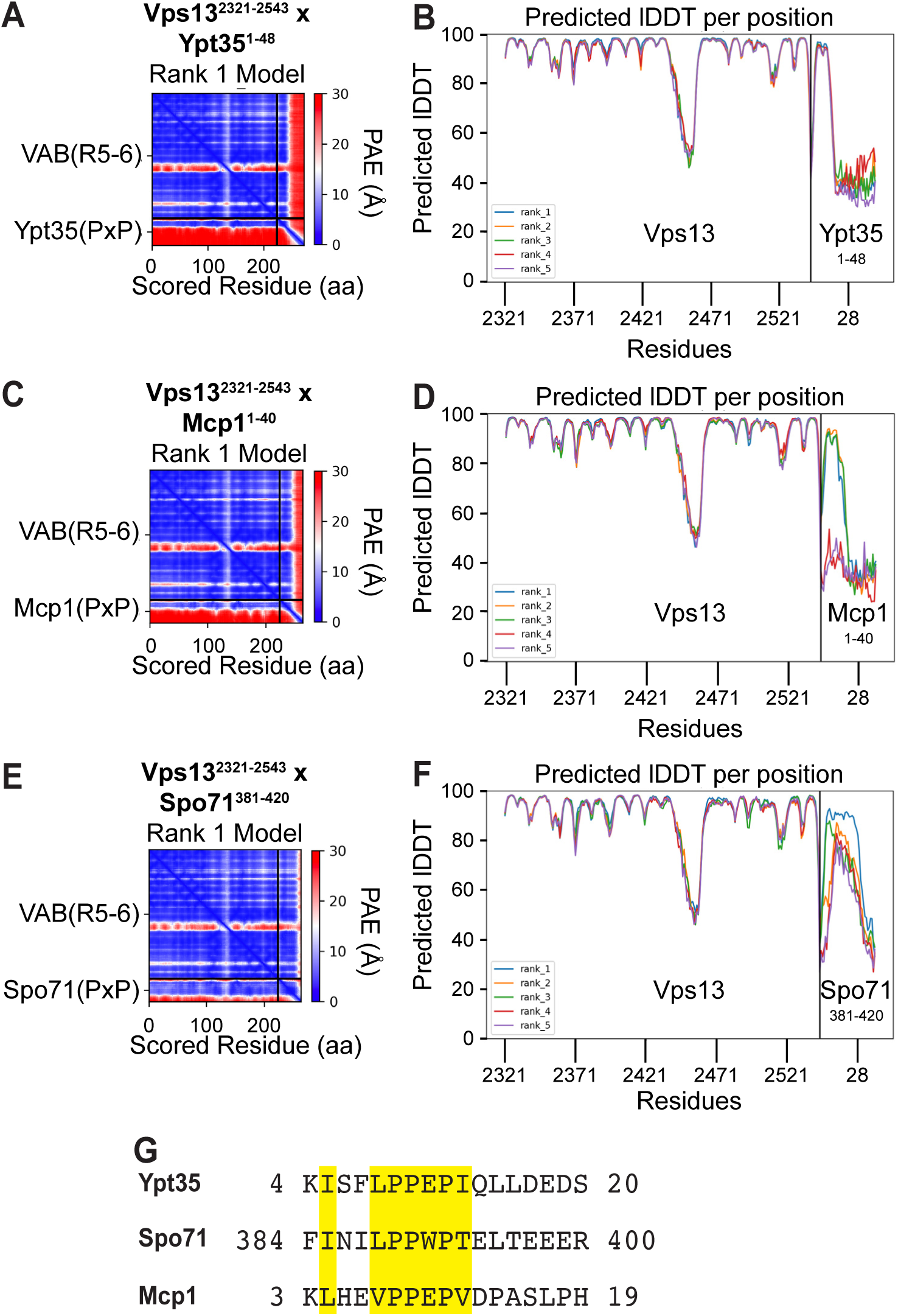
Specifications of AlphaFold and ColabFold models. (A-F) Predicted aligned error (pAE) plots (left panels) and per-residue pLDDT scores (right panels) for the ColabFold-predicted complexes shown in Figure 1 consisting of the Vps13 VAB domain bound to the PxP motif of (A and B) Ypt35^1-48^ (C and D) Mcp1^1-40^ (E and F) Spo71^381-420^. (G) Alignment of the PxP motifs of Ypt35, Spo71, and Mcp1 of *S. cerevisiae* with positions that consistently interact with VAB domain repeats 5-6 highlighted in yellow.

**Supplemental Figure 2.**
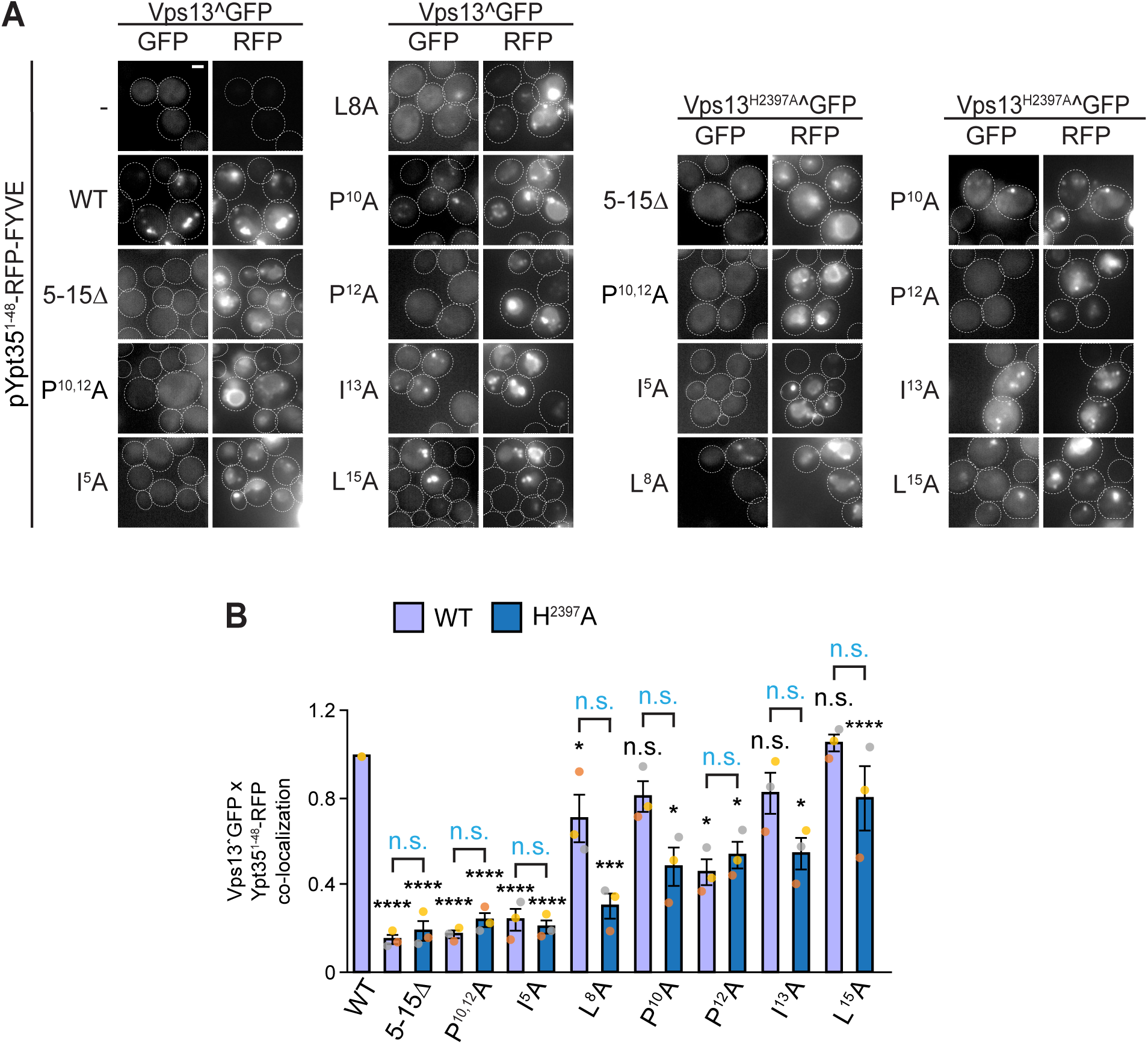
Effect of combining the Vps13 H^2397^A mutation with mutations in consensus residues of the Ypt35 PxP motif. (A) Recruitment of plasmid-expressed wildtype Vps13^GFP or Vps13^H2397A^^GFP to endosomes by chimeric adaptor proteins containing the PxP motif of Ypt35, or the indicated mutant versions, fused to the RFP-FYVE localization module. Plasmids are expressed in *ypt35*Δ yeast cells. Bar, 2µm. (B) Automated quantitation of Vps13^GFP co-localization with the Ypt35^1-48^-RFP-FYVE PxP chimera for the indicated mutant combinations, relative to total number of RFP puncta per sample. Values are relative to WT. n=3, ≥311 cells/strain/replicate. Statistical comparisons to WT are coloured black. Pairwise comparisons between WT and H^2397^A mutations in combination with the indicated Ypt35 PxP motif mutations are coloured in light blue. Two-way ANOVA test with Tukey correction, P*<0.05, P***<0.001, P****<0.0001, n.s. = not significant. Error bars = S.E.M.

**Supplemental Figure 3.**
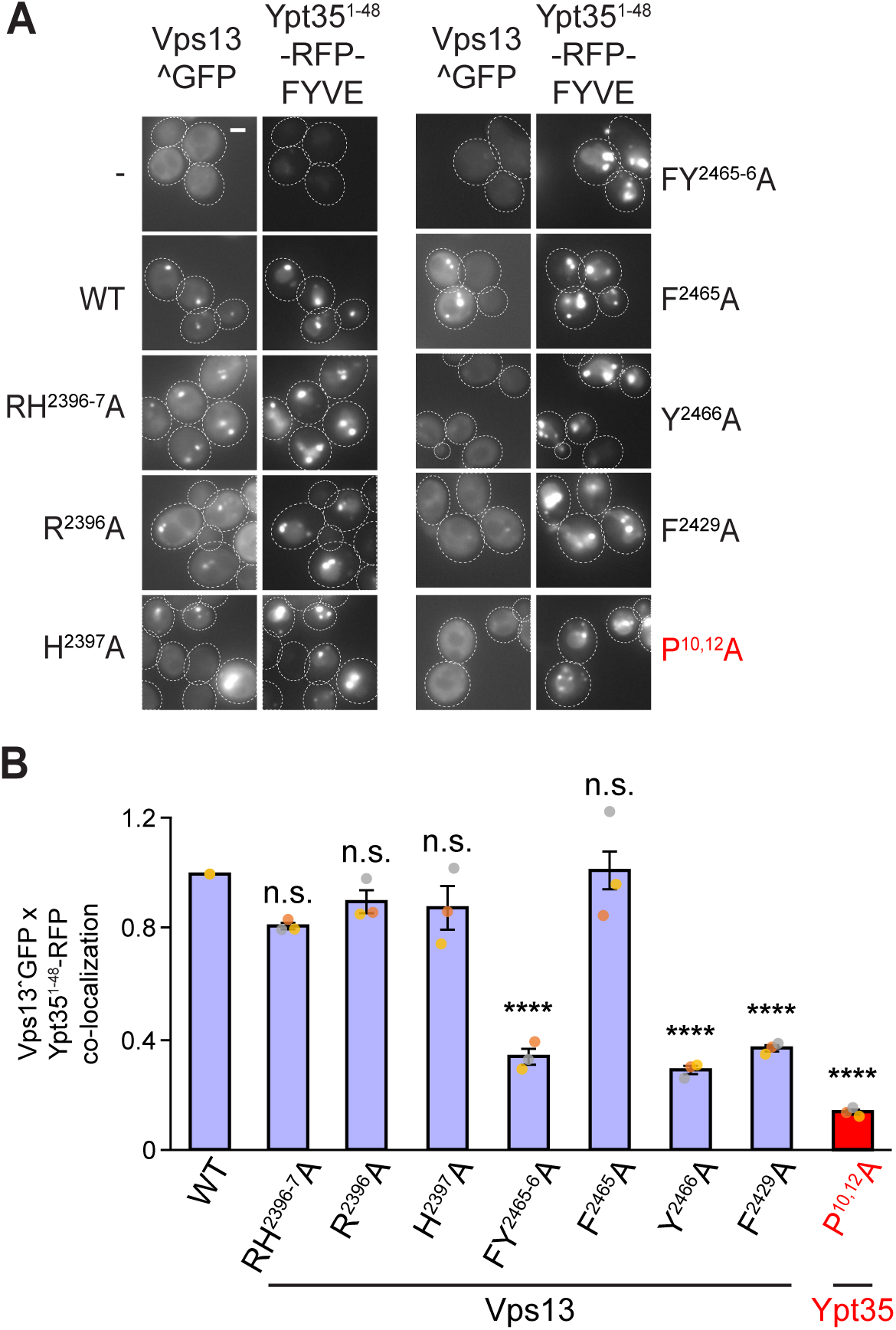
Additional VAB mutations disrupt recruitment by PxP adaptors. (A) Co-localization of wildtype and mutant Vps13^GFP with either the wild type or the P^10,12^A mutant form of the Ypt35 PxP motif fused to RFP-FYVE. Bar, 2µm. (B) Quantitation of co-localizing puncta, relative to total number of RFP puncta per sample, from (A), normalized to WT levels, n=3, ≥188 cells/strain/sample. One-way ANOVA test with Dunnett correction, P****<0.0001, n.s. = not significant. Individual data points are coloured by replicate. Error bars = S.E.M.

## REFERENCES

Adlakha, J, Hong, Z, Li, P, and Reinisch, KM (2022). Structural and biochemical insights into lipid transport by VPS13 proteins. J Cell Biol 221, e202202030.

Ashkenazy, H, Abadi, S, Martz, E, Chay, O, Mayrose, I, Pupko, T, and Ben-Tal, N (2016). ConSurf 2016: an improved methodology to estimate and visualize evolutionary conservation in macromolecules. Nucleic Acids Res 44, W344–W350.

Bean, BDM, Dziurdzik, SK, Kolehmainen, KL, Fowler, CMS, Kwong, WK, Grad, LI, Davey, M, Schluter, C, and Conibear, E (2018). Competitive organelle-specific adaptors recruit Vps13 to membrane contact sites. J Cell Biol 217, 3593–3607.

Chen, VB, Arendall 3rd, WB, Headd, JJ, Keedy, DA, Immormino, RM, Kapral, GJ, Murray, LW, Richardson, JS, and Richardson, DC (2010). MolProbity: all-atom structure validation for macromolecular crystallography. Acta Crystallogr D Biol Crystallogr 66, 12–21.

Conibear, E, and Stevens, TH (1998). Multiple sorting pathways between the late Golgi and the vacuole in yeast. Biochim Biophys Acta 1404, 211–230.

Covill-Cooke, C, Kwizera, B, López-Doménech, G, Thompson, CO, Cheung, NJ, Cerezo, E, Peterka, M, Kittler, JT, and Kornmann, B (2024). Shared structural features of Miro binding control mitochondrial homeostasis. EMBO J 43, 595–614.

Dall’Armellina, F, Stagi, M, and Swan, LE (2023). In silico modeling human VPS13 proteins associated with donor and target membranes suggests lipid transfer mechanisms. Proteins 91, 439–455.

De, M, Oleskie, AN, Ayyash, M, Dutta, S, Mancour, L, Abazeed, ME, Brace, EJ, Skiniotis, G, and Fuller, RS (2017). The Vps13p-Cdc31p complex is directly required for TGN late endosome transport and TGN homotypic fusion. J Cell Biol 216, 425–439.

Du, Y, Fan, X, Song, C, Chang, W, Xiong, J, Deng, L, and Ji, WK (2024). Sec23IP recruits VPS13B/COH1 to ER exit site-Golgi interface for tubular ERGIC formation. J Cell Biol 223, e202402083.

Dziurdzik, SK, Bean, BDM, and Conibear, E (2018). An interorganellar bidding war: Vps13 localization by competitive organelle-specific adaptors. Contact 1.

Dziurdzik, SK, Bean, BDM, Davey, M, and Conibear, E (2020). A VPS13D spastic ataxia mutation disrupts the conserved adaptor-binding site in yeast VPS13. Hum Mol Genet 29, 635–648.

Dziurdzik, SK, and Conibear, E (2021). The Vps13 family of lipid transporters and its role at membrane contact sites. Int J Mol Sci 22, 2905.

Gao, J, Franzkoch, R, Rocha-Roa, C, Psathaki, OE, Hensel, M, Vanni, S, and Ungermann, C (2025). Any1 is a phospholipid scramblase involved in endosome biogenesis. J Cell Biol 224, e202410013.

Guillen-Samander, A, Leonzino, M, Hanna IV, MG, Tang, N, Shen, H, and De Camilli, P (2021). VPS13D bridges the ER to mitochondria and peroxisomes via Miro. J Cell Biol 220, e202010004.

Hanna, M, Guillén-Samander, A, and De Camilli, P (2023). RBG Motif Bridge-Like Lipid Transport Proteins: Structure, Functions, and Open Questions. Annu Rev Cell Dev Biol 39, 409–434.

Henne, WM, Buchkovich, NJ, and Emr, SD (2011). The ESCRT pathway. Dev Cell 21, 77–91.

Jackson, CL, Walch, L, and Verbavatz, JM (2016). Lipids and their trafficking: an integral part of cellular organization. Dev Cell 39, 139–153.

Janke, C et al. (2004). A versatile toolbox for PCR-based tagging of yeast genes: new fluorescent proteins, more markers and promoter substitution cassettes. Yeast 21, 947–962.

John Peter, AT, Herrmann, B, Antunes, D, Rapaport, D, Dimmer, K, and Kornmann, B (2017). Vps13-Mcp1 interact at vacuole-mitochondria interfaces and bypass ER-mitochondria contact sites. J Cell Biol 216, 3219–3229.

Jumper, J et al. (2021). Highly accurate protein structure prediction with AlphaFold. Nature 596, 583–589.

Kolakowski, D, Kaminska, J, and Zoladek, T (2020). The binding of the APT1 domains to phosphoinositides is regulated by metal ions in vitro. Biochim Biophys Acta - Biomembr 1862, 183349.

Krissinel, E, and Henrick, K (2007). Inference of macromolecular assemblies from crystalline state. J Mol Biol 372, 774–797.

Kumar, N, Leonzino, M, Hancock-Cerutti, W, Horenkamp, FA, Li, P, Lees, JA, Wheeler, H, Reinisch, KM, and De Camilli, P (2018). VPS13A and VPS13C are lipid transport proteins differentially localized at ER contact sites. J Cell Biol 217, 3625–3639.

Lamprecht, MR, Sabatini, DM, and Carpenter, AE (2007). CellProfiler: free, versatile software for automated biological image analysis. Biotechniques 42, 71–75.

Lang, AB, John Peter, AT, Walter, P, and Kornmann, B (2015). ER-mitochondrial junctions can be bypassed by dominant mutations in the endosomal protein Vps13. J Cell Biol 210, 883–890.

Leonzino, M, Reinisch, KM, and De Camilli, P (2021). Insights into VPS13 properties and function reveal a new mechanism of eukaryotic lipid transport. Biochim Biophys Acta Mol Cell Biol Lipids 1866, 159003.

Li, P, Lees, JA, Lusk, CP, and Reinisch, KM (2020). Cryo-EM reconstruction of a VPS13 fragment reveals a long groove to channel lipids between membranes. J Cell Biol 219, e202001161.

Longtine, MS, McKenzie, AI, Demarini, DJ, Shah, NG, Wach, A, Brachat, A, Philippsen, P, and Pringle, JR (1998). Additional modules for versatile and economical PCR-based gene deletion and modification in Saccharomyces cerevisiae. Yeast 14, 953–961.

Mirdita, M, Schutze, K, Moriwaki, Y, Heo, L, Ovchinnikov, S, and Steinegger, M (2022). ColabFold: making protein folding accessible to all. Nat Methods 19, 679–682.

Neuman, SD, Levine, TP, and Bashirullah, A (2022). A novel superfamily of bridge-like lipid transfer proteins. Trends Cell Biol 32, 962–974.

Park, JS, Hollingsworth, NM, and Neiman, AM (2021). Genetic dissection of Vps13 regulation in yeast using disease mutations from human orthologs. Int J Mol Sci 22, 6200.

Park, JS, Hu, Y, Hollingsworth, NM, Miltenberger-Miltenyi, G, and Neiman, AM (2022). Interaction between VPS13A and the XK scramblase is important for VPS13A function in humans. J Cell Sci 135, jcs260227.

Park, JS, and Neiman, AM (2012). VPS13 regulates membrane morphogenesis during sporulation in Saccharomyces cerevisiae. J Cell Sci 125, 3004–3011.

Park, JS, and Neiman, AM (2020). XK is a partner for VPS13A: a molecular link between chorea-acanthocytosis and McLeod syndrome. Mol Biol Cell 31, 2425–2436.

Park, JS, Okumura, Y, Tachikawa, H, and Neiman, AM (2013). SPO71 encodes a developmental stage-specific partner for Vps13 in Saccharomyces cerevisiae. Eukaryot Cell 12, 1530–1537.

Schneider, CA, Rasband, WS, and Eliceiri, KW (2012). NIH image to ImageJ. Nat Methods 9, 671– 675.

Suzuki, SW, West, M, Zhang, Y, Fan, JS, Roberts, RT, Odorizzi, G, and Emr, S (2024). A role for Vps13-mediated lipid transfer at the ER-endosome contact site in ESCRT-mediated sorting. J Cell Biol 223, e202307094.

Tornero-Écija, AR, Zapata-Del-Baño, A, Antón-Esteban, L, Vincent, O, and Escalante, R (2023). The association of lipid transfer protein VPS13A with endosomes is mediated by sorting nexin SNX5. Life Sci Alliance 6, e202201852.

Wang, X et al. (2025). The bridge-like lipid transport protein VPS13C/PARK23 mediates ER-lysosome contacts following lysosome damage. Nat Cell Biol 27, 776–789.

Waterhouse, AM, Procter, JB, Martin, DMA, Clamp, M, and Barton, GJ (2009). Jalview Version 2 - A multiple sequence alignment editor and analysis workbench. Bioinformatics 25, 1189–1191.

Wong, LH, Gatta, AT, and Levine, TP (2019). Lipid transfer proteins: the lipid commute via shuttles, bridges and tubes. Nat Rev Mol Cell Biol 20, 85–101.

